# Assessing the quality of generative artificial intelligence for science communication in environmental research

**DOI:** 10.1101/2024.11.11.623072

**Authors:** David Worden, Daniel Richards

## Abstract

The adoption of Generative Artificial Intelligence (GenAI) tools is drastically changing the way that researchers work. While debate on the quality of GenAI outputs continues, there is optimism that GenAI may help human experts to address the most significant environmental challenges facing society. No previous research has quantitatively assessed the quality of GenAI outputs intended to inform environmental management decisions. Here we surveyed 98 environmental scientists and used their expertise to assess the quality of human and GenAI content relevant to their discipline. We analysed the quality and relative preference between human and GenAI content across three use cases in environmental science outreach and communication. Our results indicate that the GenAI content was generally deemed adequate in quality by human experts, with an average of 82% of respondents indicating a quality of “adequate” or better across the three use cases. Respondents exhibited strong preferences for GenAI over human-only content when using GenAI imageery of future park management scenarios. For the use cases of generating a wetland planting guide and answering a question about invasive species management, preferences were heterogeneous amongst respondents. Our findings raise substantive questions about GenAI content as a complement to human expertise when research is transferred to public audiences.

## Introduction

The rapid adoption of Generative Artificial Intelligence (GenAI) tools over the past two years is drastically changing the way that researchers work (Bakker and Traniello, 2024; Van Dis et al., 2023; Zhu et al., 2023). With widespread adoption, concerns have grown around ethical uses, mis- and dis-information, high energy and water usage, data privacy and sovereignty, and the potential impact to job security (Schlagwein and Willocks, 2023; Hacker et al., 2023; Li et al., 2024; Eloundou et al., 2023; Lucchi, 2023; Hicks et al., 2024). In contrast to these emerging concerns, proponents have expressed optimism about the potential of GenAI to accelerate progress on some of societies biggest issues such as climate change and environmental management (Biswas, 2023; Zhu et al., 2023; Rane et al., 2024).

GenAI has been identified as a potential solution for environmental science communication, to help bridge the gap between scientists and practitioners, and rapidly disseminate advice and information to support decision-making (Rillig et al. 2023; Larosa et al. 2023; Vaghefi et al. 2023; Richards et al. 2024). Science outreach represents a necessary but often difficult and time-consuming task for many researchers (Andrews et al., 2005; Mitton et al., 2007). Scientists often struggle to provide policy relevant outputs from complicated analysis and GenAI may help to provide a user-friendly way of generating public-oriented information derived from scientific research in a cost-effective and scalable way.

For GenAI to be widely adopted for science communication it must be useful. This will require that the quality of GenAI content is sufficient that more time is spent on accelerating scientific progress rather than on editing or supervising GenAI outputs. Science communication may be particularly well-positioned to benefit from GenAI as researchers are often constrained when it comes to outreach and communication tasks as opposed to other elements of their work (Andrews et al., 2005; Mitton et al., 2007). Dwivedi et al. (2023) provided the first major review of the use GenAI for research purposes across a wide range of disciplines with 43 authors’ analyses of ChatGPT. While the authors disagreed on the role of regulation in GenAI adoption, they unanimously agreed that GenAI will likely offer significant productivity gains and be transformative in a range of fields. Many studies have documented the ability of GenAI – most commenting on ChatGPT – to be able to assist in research tasks. Merow et al. (2023) note the role of ChatGPT in improving productivity in coding tasks; Zhu et al. (2023) and Huang and Tan (2023) note the role of ChatGPT in improving writing and literature reviews; Korinek (2023), Agathokleous et al (2023), and Boyko et al (2023) highlight the value of ChatGPT for ideation and brainstorming. In more specific use cases, Juhasz et al (2023) demonstrate the multimodal value of ChatGPT to supplement geospatial data by prompting ChatGPT with street-view imagery and Richards et al (2024) examine the benefits of GenAI for reporting on agricultural environmental performance, interactive biodiversity-enhancing garden design, and image generation for future landscape scenarios.

A growing body of research takes a more sceptical approach to GenAI and notes the significant and unresolved concerns present among academics when it comes to GenAI adoption. Issues of copyright, hallucinations and information inaccuracies, intellectual property, and plagiarism feature prominently in multiple articles on GenAI adoption in academia (Alkassi and McFarlane, 2023; Lucchi, 2023; Bakker and Traniello, 2024; Thorp, 2023; Hicks et al., 2024). Several authors have called for more specific and stringent guidelines or regulations on appropriate GenAI use (Schlagwein and Willcocks, 2023; Birhane et al., 2023; Hacker et al, 2023) while studies such as Ferrara (2024) point out the potential flood of information manipulation that may follow widespread GenAI adoption.

In more pragmatic terms, Taddeo et al. (2021) and Bender et al. (2021) provided an important perspective on the environmental cost of training and operating LLMs prior to their widespread adoption. While they note that AI could play an important role in addressing the climate emergency, the energy costs of providing this technology are substantial and could increase quickly. Since the widespread adoption of GenAI in 2023 other studies have confirmed this prediction by Taddeo et al. (2021). The energy and carbon costs of AI deployments are significant and require careful consideration although further research will be needed to examine these impacts over time (Luccioni et al., 2024).

From an epistemological perspective, Messeri & Crockett (2024) examine what knowing and knowledge generation might be reduced to with reliance on GenAI. They describe ‘monocultures of knowing’ whereby the ability of researchers to make innovative discoveries may be significantly diminished by funnelling research into the narrow scope of tasks suitable for AI tools and their training data. Van Dis et al. (2023) note that ChatGPT is undeniably a ‘game-changer’ for science but outline guiding principles for its use which include maintaining human verification, developing rules for accountability, making LLMs open and transparent, embracing the benefits of AI, and furthering the debate and discussion on the issue.

Few studies unequivocally support or oppose the use of GenAI in research with most studies highlighting both potential benefits and pitfalls. In large part, the concerns mentioned above remain unresolved. Fewer studies have sought to examine the role that GenAI might play in scientific communication. Alvarez et al. (2024) argue there are several benefits of GenAI for science communication such as in the ability to offer dialogical understanding on a topic or in translational outputs but that the risks of perpetuating biases and missing critical nuances – especially on sensitive research topics – cannot be ignored. Schäfer (2023) notes that GenAI may allow researchers to provide dialogical science communication at scale which could revolutionize science communication but that this also represents a double-edged sword whereby it could lead to ‘wrongness at scale’ through inaccuracies and hallucinations perpetuating misinformation in science. Biyela et al. (2024) provide a brief but thorough overview of benefits and risks in using GenAI for science communication while noting that GenAI reflects back its training data in responses so the tool could be a useful way for science communicators to identify topics that have been poorly communicated about in the past.

Lastly, on ecological research, Rillig et al. (2023) argue that communication among researchers and with the wider public are some of the biggest potential advantages to GenAI although misinformation and the environmental impact of GenAI must also be carefully considered. Richards et al. (2024) highlight the strength of GenAI in three use cases of content for public audiences including reports for the environmental performance of farms, garden designs for native biodiversity, and using image generation to assess the trade-offs between economic and biodiversity benefits. Lastly, Vaghefi et al. (2023) examined the quality of ChatGPT outputs when training data was supplemented with the Sixth Assessment Report of the Intergovernmental Panel on Climate Change (IPCC) finding that, with appropriate prompting and supplementation of the training data, more accurate and helpful responses were provided by the chatbot.

In this study, we provide initial evidence of the quality of GenAI-human content when compared with human-only content in environmental science communication use cases. We distinguish between human-only (no use of GenAI) and GenAI-human (GenAI from this point onwards) to highlight that GenAI content does not exist without human input via prompting and that the more accurate distinction is not over whether a human researcher or science communicator is involved but rather whether GenAI tools have been used to create content or not. Rather than focusing on core research activities such as data collection or analysis (Dwivedi et al., 2023; Merow et al., 2023; Zhu et a., 2023), we examined use cases in the realm of public outreach and communication as this represents a critical frontier in monitoring potential misinformation and mis-alignment of GenAI outputs with public understanding of science and knowledge discovery. Using a blind experiment administered through a survey, we aimed to examine the quality of human-only and GenAI content as expressed by the preference of expert researchers and support staff. We assessed quality across three different environmental science communication use cases; (1) an infographic wetland restoration guide, (2) a response to a public question following a presentation about invasive species, and (3) an image representing a proposed urban park scenario. These use cases were delivered through three GenAI tools, respectively *ChatGPT* (specifically *GPT-4*), Google’s *Bard* (now *Gemini*), and *Midjourney*. For each use case we analysed differences in the perceived quality of the human-only and GenAI-content, and the participant preference for the human-only or GenAI-human content.

## Materials and methods

### Survey Design

We distributed the survey to staff at an environmental and ecological research institute in Aotearoa New Zealand over a three-week period in September 2023. The survey was distributed online through Qualtrics via an anonymous link. The survey underwent social ethics review and approval at the host institution and respondents participated with informed consent and were able to cease their participation at any time. At the time of survey distribution, only 39% of respondents had used generative artificial intelligence tools for research purposes and of that sub-sample, over 97% had only used *ChatGPT* reflecting its dominance in the GenAI space. We received 98 responses to the survey and 57% were from individuals working in research roles and 43% were comprised of support, technician, student, or managerial roles.

Respondents were shown three pairs of content for each use case in a blind experiment (i.e., content was displayed in a randomised order and not labelled to allow for distinguishing between GenAI or human-only). For a breakdown of the summary statistics for the respondents see Table 1 below. The survey utilised a within-subjects survey design, to address researcher heterogeneity, which was of greater concern given our sample size. By utilising this design, each respondent served as their own control when comparing human-only versus GenAI content. A potential concern in within-subject surveys is that ordering of the experimental elements can lead to biases as respondents see both treatment and control states (Charness et al., 2012). To mitigate this issue, we randomised the order with which elements were presented to respondents as well as whether elements were presented on the left (top) or right (bottom) side of the screen in comparisons. With three use cases and the ordering randomised, this led to a total of twelve survey versions. Crucially, with this randomisation and no labelling of whether content was authored by GenAI or a human, respondents were not able to know with certainty whether the content they were seeing was made with GenAI tools or not. Respondents were able to click on figures in the survey to open them in more detail in a separate browser tab if they wished to inspect details of the content more closely such as with the visual elements in the first and third use cases.

**Table 1.** Summary Statistics for survey respondents.

| **Variable** | **N** | **Mean** | **Std. dev.** |
|---|---|---|---|
| Has used GenAI (1 = Yes) | 96 | 0.385 | 0.489 |
| Has used ChatGPT (1 = Yes) | 37 | 0.973 | 0.164 |
| Role – Researcher (1 = Yes) | 95 | 0.568 | 0.498 |
| Role - Technician / Student / Extension (1 = Yes) | 95 | 0.274 | 0.448 |
| Role - Support / Management (1 = Yes) | 95 | 0.158 | 0.367 |
| Discipline – Ecology (1 = Yes) | 82 | 0.439 | 0.499 |
| Discipline - Social science (1 = Yes) | 82 | 0.122 | 0.329 |
| Discipline – Others (1 = Yes) | 82 | 0.439 | 0.499 |
| Highest degree earned less than 5 years ago (1 = Yes) | 97 | 0.268 | 0.445 |
| Highest degree earned 5 to 20 years ago (1 = Yes) | 97 | 0.495 | 0.503 |
| Highest degree earned more than 20 years ago (1 = Yes) | 97 | 0.237 | 0.428 |

We sought to assess the quality of GenAI tools for scientific communication purposes. We chose three different types of outreach content and used a different GenAI tool for each of them. An overview of the three science communication use cases is outlined in Table 2 and expanded upon in the next section. GenAI tools are developing rapidly and have transformed significantly since their widespread public availability in 2023. The way in which GenAI tools can be used can vary significantly with more technical users often utilising an API for access or providing their own supplemental training data to ‘weight’ a LLM towards data that they want to prioritise in content generation. We utilised the default ‘off-the-shelf’ versions of each of the GenAI tools in order to provide an assessment of the baseline and most widely used versions of these tools. These versions represent the lowest barriers to entry requiring relatively little technical knowledge to be able to prompt for content.

**Table 2.** Overview of Science Communication Use Cases.

| **Use Case** | **Communication task** | **GenAI tool used** | **Original human-only content reference** |
|---|---|---|---|
| One (UC1) | A wetland planting / restoration guide with details on what species to plant as well as a planting guide and management of weeds and pests. | *ChatGPT* (*GPT-4*) | Tanner et al., 2022; Waikato Regional Council, 2018 |
| Two (UC2) | Responding to a text-based public query about a presentation on invasive species management. | *Bard (now known as Gemini)* | Manaaki Whenua, 2020a; Manaaki Whenua 2020b |
| Three (UC3) | Image generation to depict an urban park management scenario | *Midjourney* | Barron et al., 2023 |

### Use Case One: Wetland Planting / Restoration Guide

In the first use case we provided a one-page text and image summary to respondents which conveyed information about a wetland planting and restoration guide. The human-only content version was an amalgamation of two original pieces of human generated content. This approach was chosen to help alleviate concerns that survey respondents may be familiar with a document on this topic as several of the respondents in our study had expertise in wetland ecology. The first piece of human content was a constructed wetland practitioner guide produced by DairyNZ (an industry organisation for dairy farmers in Aotearoa New Zealand) and NIWA (a crown research institute in Aotearoa New Zealand specialising in freshwater, climate and ocean research) which provided the information on planting zones and ideal species (Tanner et al., 2022). The second piece of content was a planting guide authored by the Waikato Regional Council (a regional council of Aotearoa New Zealand) which was the basis of the guide headings for preparing a planting plan, timing, site preparation, planting and weeds and pests (Waikato Regional Council, 2018). These two sources of information were edited and combined into one infographic (Figure S1), which was displayed to survey respondents as the human-only content option.

Using this same template, we then used *ChatGPT* (*GPT-4*) to populate each section with content (OpenAI, 2023). The template was populated with content by prompting for each relevant heading individually but in the same continuous dialogue with *ChatGPT* (*GPT-4*). In this continuous dialogue it was possible to reiterate that *ChatGPT* should refer back to its own content for consistency and to the context of Aotearoa New Zealand. The prompts used to generate content for each section are provided in the Supplementary Methods. The GenAI infographic shown to respondents for use case one is presented in Figure S2.

### Use Case Two Responding to a Query from the Public

The second science communication use case we examined was responding to a question from the public ex-post an online presentation. The human-only content we presented to survey respondents was a text question and answer relating to an online presentation about invasive species (wilding pines – *Pinus radiata*) management in Aotearoa New Zealand. The presentation was available on *YouTube* (Manaaki Whenua, 2020a) and the presenters answered additional questions after the presentation and provided a text Q&A document (Manaaki Whenua, 2020b). This afforded us an example to be able to prompt a GenAI chatbot to consider the context and knowledge within a *YouTube* presentation when generating its response. We used *Google*’s *Bard* (now known as *Gemini*) chatbot for this task because it was able to respond to links to *YouTube* and provide feedback on questions relating to the *YouTube* video (Google, 2023). This task reflected an example where rather than a science communicator responding in real time, they were able to take time to answer a question with a text-response at a later date which, could be, a suitable example where GenAI could be used to fill this role. The human-only and GenAI content are presented in Table S1 and S2.

The human-only content was edited slightly for grammar and typographical errors to ensure it would not be assessed on the basis of those elements. For prompting the GenAI chatbot *Bard*, we first prompted *Bard* to ask if it was familiar with the *YouTube* video. Once it responded indicating it could access that content for reference, we asked it to roleplay as if it was one of the presenters when it came to answering the question. The full prompts provided to *Bard* are provided in the Supplementary Methods.

Following assessment of each use case, respondents had the option of providing qualitative feedback on both the human-only and GenAI content. Several respondents noted that the GenAI content for this use case was verbose and overly optimistic as compared to the human-only content which is much more succinct and to the point of the question. Additionally, the GenAI response mentions specific details such as naming two companies that use oils extracted from wilding conifers. At the time of writing, it is unclear whether the specific named companies exist although it does appear that similarly named companies are operational. The human science communicator that answered this question also referenced oils from wilding conifers but chose to not specifically name a company, which likely reflects the nuances of professionalism that may be lost on a chatbot.

### Use Case Three Landscape Image to Reflect Habitat Restoration

In the third science communication task we examined the role of imagery to convey ecological research and concepts. This task presented several challenges as we needed to find a recent and relevant image that also had a detailed text description from the creators of the image which would allow a fair comparison given the intentions of the original human-only image and without imparting our own perspective into describing the elements that made up an image. We used the text-to-image GenAI tool *Midjourney* for this task (Midjourney, 2023). Once again, we wanted to find a recent example of content that was unlikely to be recognised by our survey respondents that had expertise in this subject matter. For this reason, we used an image based in a context outside of Aotearoa New Zealand (based in Canada) but provided survey respondents with this context when they were assessing the images. The human-only image we used was from a recent article by Barron et al. (2023) which examined urban park design in a case study located in Vancouver, Canada (Figure S3). In the supplemental material for Barron et al. (2023) they provided a text description of each of the design scenarios they present in their paper which we modified to prompt *Midjourney* for image creation (Supplementary Methods). The GenAI restoration design concept is shown in Figure S4.

### Empirical method

In the survey we asked respondents to respond to the content as if they were peer-reviewing them for a colleague before wider distribution. Respondents were made aware that each pair of content would include one human-only and one GenAI example but that the ordering of those would be randomised. The respondents were asked to rate the quality of each content in a pair using a five-point Likert scale (i.e., from very poor to very good) as well as to provide a relative ranking on a five-point scale assessing which content was relatively better (i.e., from strongly preferring GenAI to strongly preferring human-only content). Respondents were also asked to assess how relevant the use case was to their specific research (from not at all relevant to very relevant).

To directly compare human-only content versus GenAI content, we also asked respondents to provide their relative preference between each pair for each use case. This additional assessment is important because respondents may have ranked the quality of both human-only and GenAI content in the same category but by asking respondents to indicate a preference we improve the robustness of assessing whether human-only or GenAI content is performing better for the three science communication use cases we examined.

Paired t-tests and Wilcoxon matched-pairs signed-rank tests were used for robustness as the discussion of whether ordinal data such as Likert-scale responses should be analysed with parametric or non-parametric tests is still subject to debate (de Winter and Dodou, 2010; Derrick and White, 2017). The results presented below use the full sample of 98 respondents. The findings are robust to both paired t-tests and Wilcoxon matched-pairs signed-rank tests as well as to whether using the full sample or just the sub-sample of respondents that indicated the content was at least somewhat relevant to their work as well as the sub-sample of only researchers (omitting technician, support, student, extension, and management roles).

For the paired t-test on quality ratings we assess whether the difference between the mean of the quality values for each pair of content are equivalent as well as performing the one-sided tests of the distribution which indicate whether there is a higher quality for either human-only or GenAI content. The non-parametric Wilcoxon matched-pairs signed-rank tests are performed similarly but assess whether a difference exists across median, as opposed to mean, values in addition to performing the one-sided tests that are necessary to establish which content is preferred if the mean or median difference is not equal to zero.

T-tests and Wilcoxon signed-rank tests are also used for assessing preferences between human-only and GenAI content but in this case the tests are used to assess whether the mean or median value is equivalent to the central moment of the Likert-scale (i.e., the mean or median is equal to three) which would reflect no preference between GenAI or human-only content. One sided-tests are also performed for the preference rankings which is necessary to determine the direction of the preference (i.e., preferring human-only or GenAI content when the mean or median is not equivalent to three).

## Results

The human-only and GenAI outputs were generally perceived as adequate in quality across the use cases (see Figure 1 below). There was no statistically significant difference in the scoring of output quality between the human-only and GenAI outputs for the first and second use cases. The only statistically significant result in assessing quality was for the third use case, in which respondents significantly rated the quality of the output that had been generated with *Midjourney* over the original image from the Barron et al. (2023) publication. Importantly, being unable to reject the null hypothesis in assessing quality in Use Case One (UC1) and Use Case Two (UC2) indicates that our sample of experts were not able to make definitive assessments about the quality of human-only content being better than GenAI content, or vice versa.

**Figure 1.**
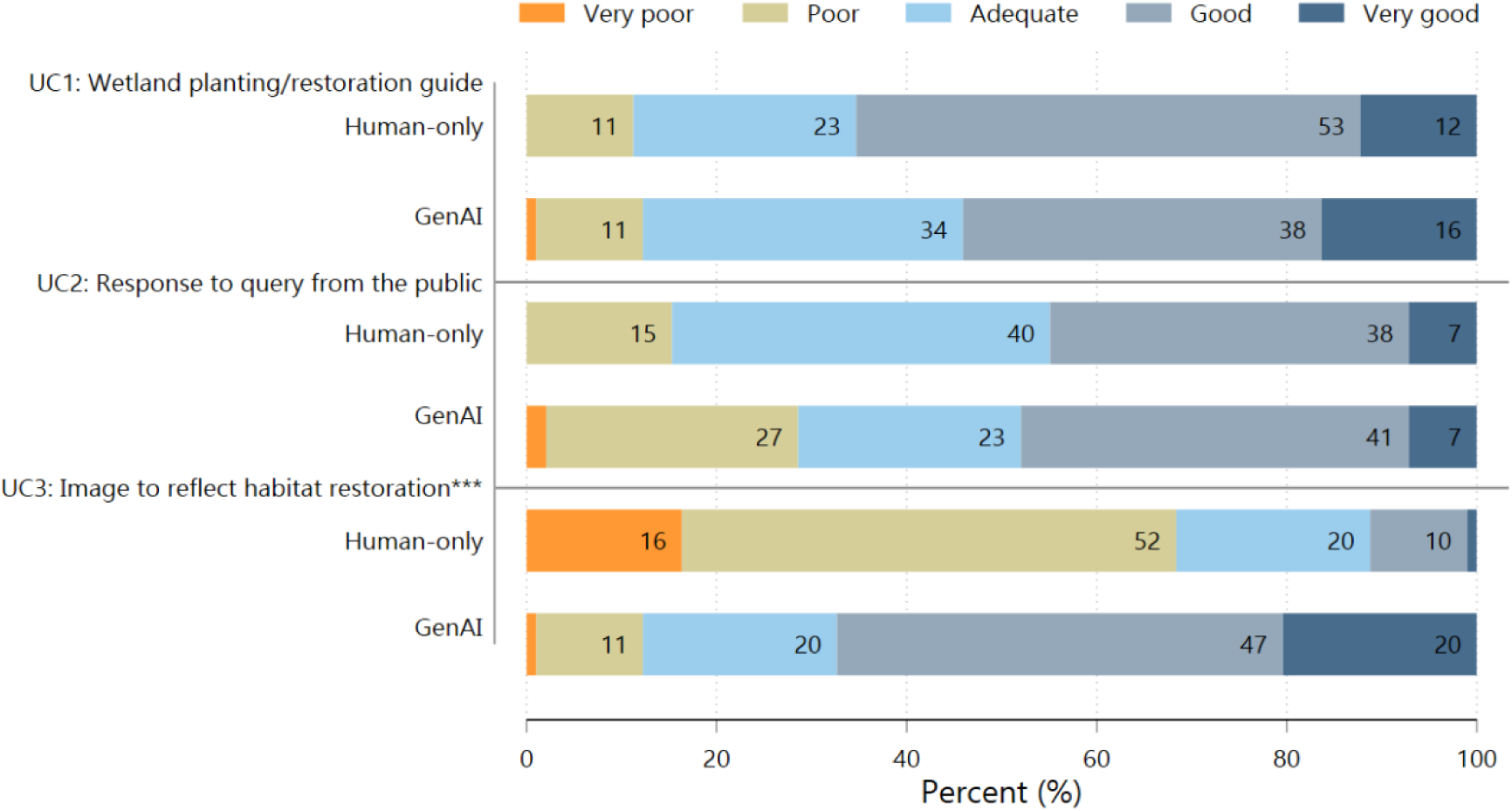
Quality of Human-only and GenAI content across three use cases. N = 98, t-test and Wilcoxon signed-rank p-values: * p<0.05, ** p<0.01, *** p<0.001 H_0_ : UC_i_ Human quality = UC_i_ GenAI quality; H_a_ : UC_i_ Human quality ≠ UC_i_ GenAI quality; H_a_: UC_i_ Human quality > UC_i_ GenAI quality; H_a_ : UC_i_ Human quality < UC_i_ GenAI quality

The Likert rankings of preferences between GenAI and human-only content were divided for UC1 and UC2, indicating diverse preferences between respondents (Figure 2). Preferences between GenAI and human-only content were more balanced in the first use case, with similar numbers of people preferring the different use cases and expressing no preference. In the second use case opinions were more divided, with fewer respondents expressing no preference. In UC3 there was a significant preference among respondents for the GenAI image over that of the human-only image of the habitat restoration in an urban park.

**Figure 2.**
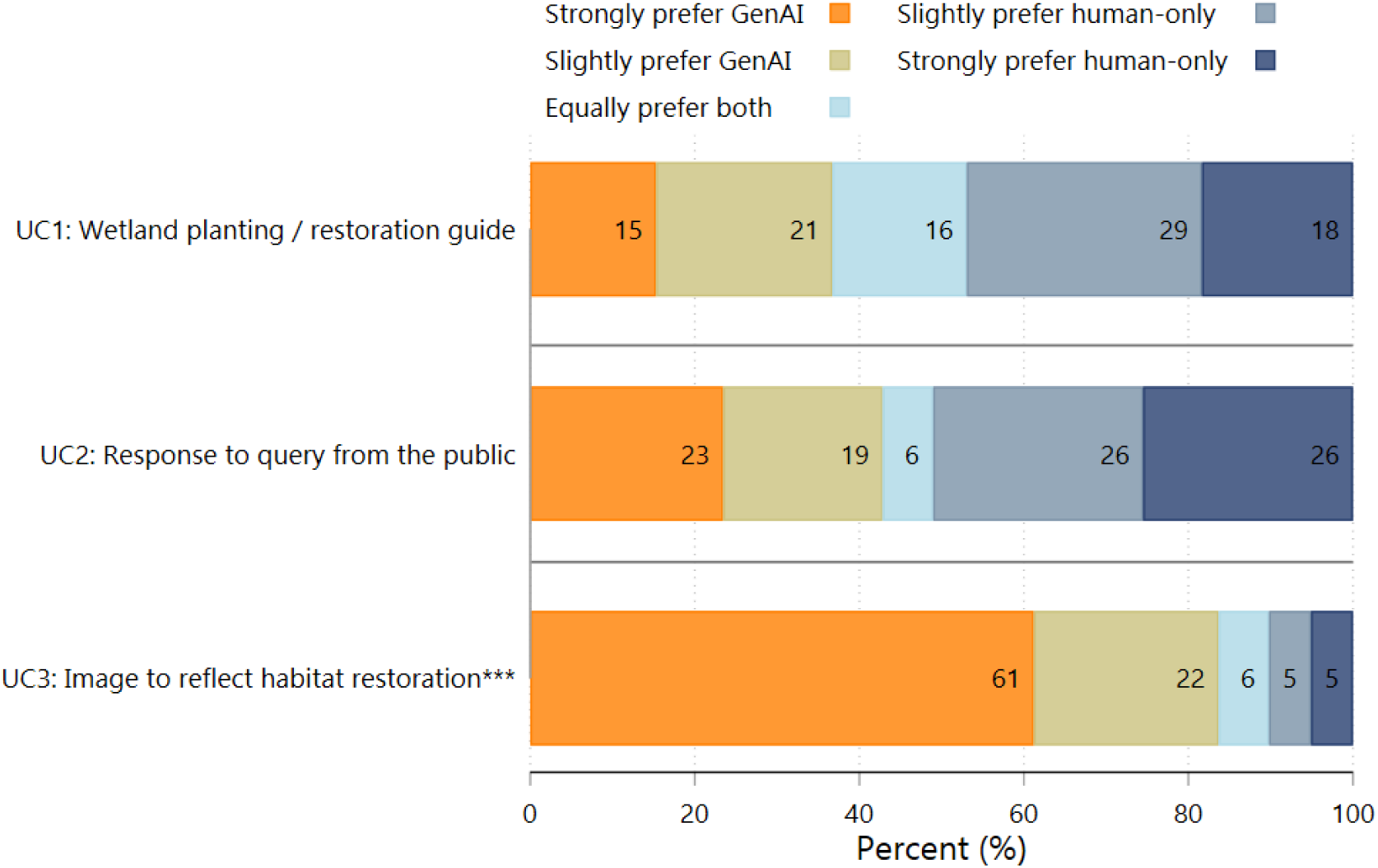
Preferences between Human-only or GenAI content across three use cases. N=98, t-test and Wilcoxon signed-rank p-values: * p<0.05, ** p<0.01, *** p<0.001 H_0_ : UC_i_ Preference = 3 (equally prefer both); H_a_ : UC_i_ Preference ≠ 3 (equally prefer both); H_a_ : UC_i_ Preference > 3 (prefer human); H_a_ : UC_i_ Preference < 3 (prefer GenAI)

## Discussion

While studies such as Alvarez et al. (2024), Schäfer (2023), Biyela et al. (2024), Rillig et al. (2023), and Richards et al. (2024) provide anecdotal evidence of the potential benefits of GenAI for science communication, ours is the first study to undertake a quantitative quality and preference assessment comparing GenAI and human-only content on scientific outreach tasks. Our results suggest that GenAI content may be comparable in terms of quality to human-only content across at least some science communication tasks. This requires careful consideration from the research and scientific outreach community. We do not intend for this result to condone a laissez-faire approach to GenAI use in research or scientific communication and we stress the importance of maintaining human oversight and accountability for the use of GenAI content, however these findings suggest that there could be significant advantages for researchers and scientists that feel constrained in their ability to communicate about their work to other disciplines or broader public audiences. With some knowledge of prompt engineering and the ability to provide oversight on outputs, researchers may be able to use GenAI tools to scale up their outreach abilities in a cost-effective way.

For the wetland planting / restoration guide and the response to a query from the public on a presentation there was no statistically significant difference in how respondents assessed quality or preference rankings between GenAI and human-only content. The summary statistics of the results presented in Figure 6 and 7 may suggest that with a larger sample size, human-only content could be rated of a higher quality and preferred in these two use cases. However, it is important to note that GenAI content is still preferred 36% and 42% of the time between use case one and use case two, respectively. In other words, GenAI content may be ‘good enough’ for certain types of scientific communication tasks especially when factoring in that 82% of our respondents thought the GenAI content was at least “adequate” on average across the three use cases. This, coupled with the time savings that GenAI use may afford researchers, suggests that it may be a worthwhile tool for accelerating and scaling up science communication.

Prompt engineering is a critical element to obtaining useful content (Richards et al. 2024). For example, in Use Case One (UC1) we provided *ChatGPT* (*GPT-4*) with a predefined structure to its responses and the context that it should be providing wetland restoration / planting suggestions in the context of Aotearoa New Zealand. Without providing this context to prompts, the output for *ChatGPT* is likely to be more general and may be unhelpful – such as recommending invasive plant species. In Use Case Two (UC2) *Google’s Bard* chatbot was able to draw from the context of a *YouTube* video to provide a text response to a question from the public.

For certain tasks, such as generating realistic landscape imagery for outreach purposes, GenAI may out-perform typical status quo approaches such as manual photo-editing or photo-collage techniques. Considering the expertise, software, and labour hours that go into high-quality photo editing, GenAI image generation tools may provide significant benefits for scientific communication tasks that rely on visual representations of the subject matter but where a high level of detail is less critical.

Our study also innovates by using three different generative artificial intelligence tools which helps to broaden the perspective on GenAI tools and uses away from the dominance of *ChatGPT*. This helps to expand the discussion beyond focusing solely on *ChatGPT*, acknowledging that certain GenAI tools such as *Midjourney* that are designed for one task – image generation – may be worthwhile for researchers and science communicators to engage with.

While inaccuracies and hallucinations of GenAI are a real challenge to their adoption for scientific communication, it is worth noting that GenAI tools may offer significant benefits to science communicators. For the first time, an easily accessible and widely available technology exists that can generate science communication content to rival the capabilities of scientists and researchers in real-world applications.

## Limitations

As with any study, ours has several limitations that require consideration. Our sample of ninety-eight respondents were drawn from the staff of one research organisation in Aotearoa New Zealand. Further research could benefit from surveying a higher number of researchers and science communicators across a global context. Second, our results represent a snapshot in time for GenAI adoption in September 2023. This snapshot was collected at a time when knowledge of GenAI and its potential applications was relatively low - most respondents had not at that point in time used GenAI for research purposes yet. Conversely, now that more people have been exposed to GenAI content and surrounding debates in the media and scientific literature, the results of the same survey conducted today may be different. As users interact more with GenAI it becomes easier for them to identify the types of content it creates. Without additional prompting or editing it is likely that a notable number of respondents would be able to accurately determine the GenAI content from that of a human.

A further challenge of our study representing this snapshot in time is that it precludes us from being able to assess the impact of new innovations in the GenAI and LLM space. In particular, the most recently released version of ChatGPT (ChatGPT-4o) allows users to interact with the chatbot via multimodal inputs such as text, images, video, and audio and be able to receive multimodal outputs as well (Edwards and Orland, 2024). In particular, the strength of video and audio innovations in GenAI tools could significantly improve the prospects of science communication by improving accessibility and enabling a wider range of interactions for communicating concepts.

Third, we examined three specific use cases out of countless potential applications for GenAI in science communication and research more broadly. It is likely that there are many use cases where GenAI performs very poorly such as in scientific communication about socially or culturally sensitive subjects and there are also likely to be cases where it performs even better than our findings suggest (Richards et al 2024).

## Conclusion

In this study we surveyed researchers and staff at a research institute to assess how GenAI content compared to human-only content in scientific communication tasks. Our findings suggest that, at least in some cases, GenAI may perform as well as human-only content when independently assessed by others in the science community. In one of the three use cases, image generation, GenAI significantly outperformed human-only content.

This study provides an important contribution to the literature on the use of GenAI tools in research and science communication as we provide some of the first evidence of quality and preference rankings when human-only and GenAI content are compared. We also provide an assessment across three GenAI tools, including *ChatGPT*, to highlight the potential value from broadening the discussion to include other GenAI innovations.

We support the conclusions of various other studies that proper oversight in the use of GenAI in research and scientific communication is necessary and stress that our results should not be seen as supporting GenAI as a substitute to human researchers or science communicators. However, our results strengthen the perspective that GenAI can significantly benefit researchers and science communicators. As we note, there are many challenges – ranging from data sovereignty and intellectual property to the energy intensiveness of GenAI models – but similar challenges have been present in past technological advancements. Disregarding the value of GenAI tools to researchers and science communicators may be even more costly.

## Acknowledgements

This work was supported by the Strategic Science Investment Funding for Crown Research Institutes from the New Zealand Ministry of Business, Innovation and Employment’s Science and Innovation Group.

## Author contributions

D.W. and D.R. wrote the main manuscript text. D.W. and D.R. designed the survey. D.W. prepared the statistical analysis and figures. D.R. produced the content for Use Case One, D.W. prepared the content for Use Case Two and Three.

## Data availability statement

Survey data presented in this study are unavailable due to confidentiality.

## Supplemental Materials

**Figure S1.**
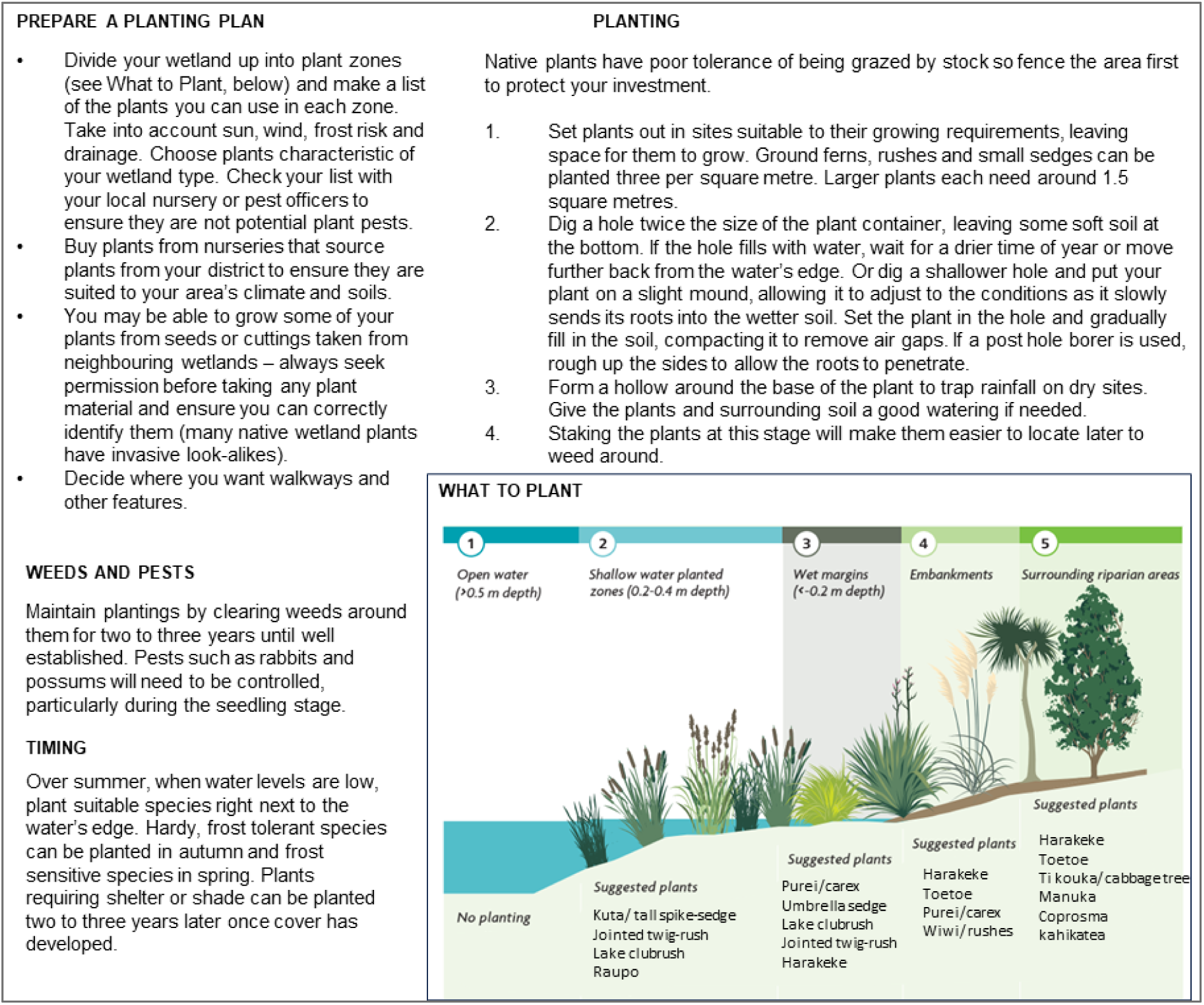
Use Case One Human-only Content

**Figure S2.**
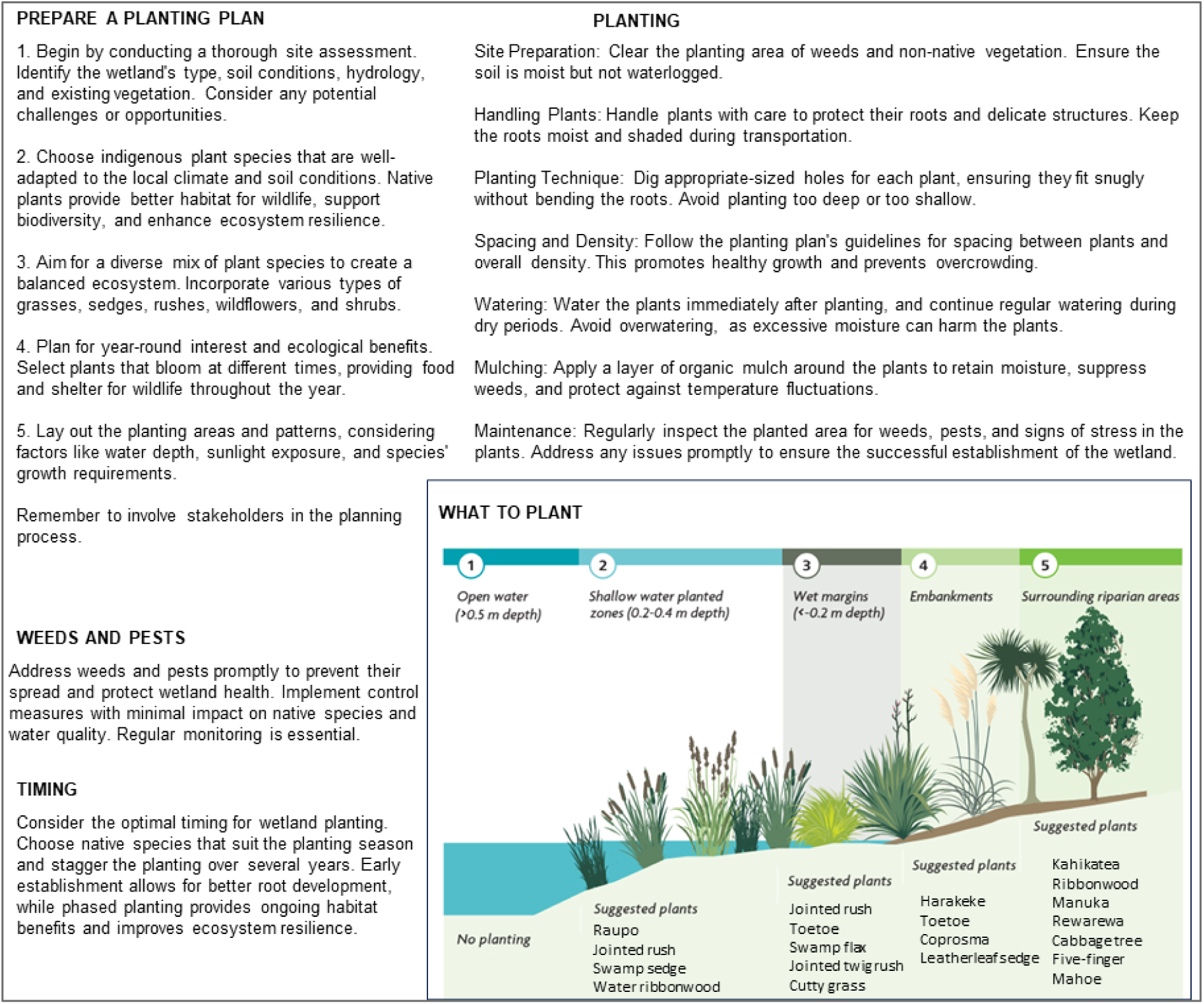
Use Case One GenAI Content

**Table S1.**
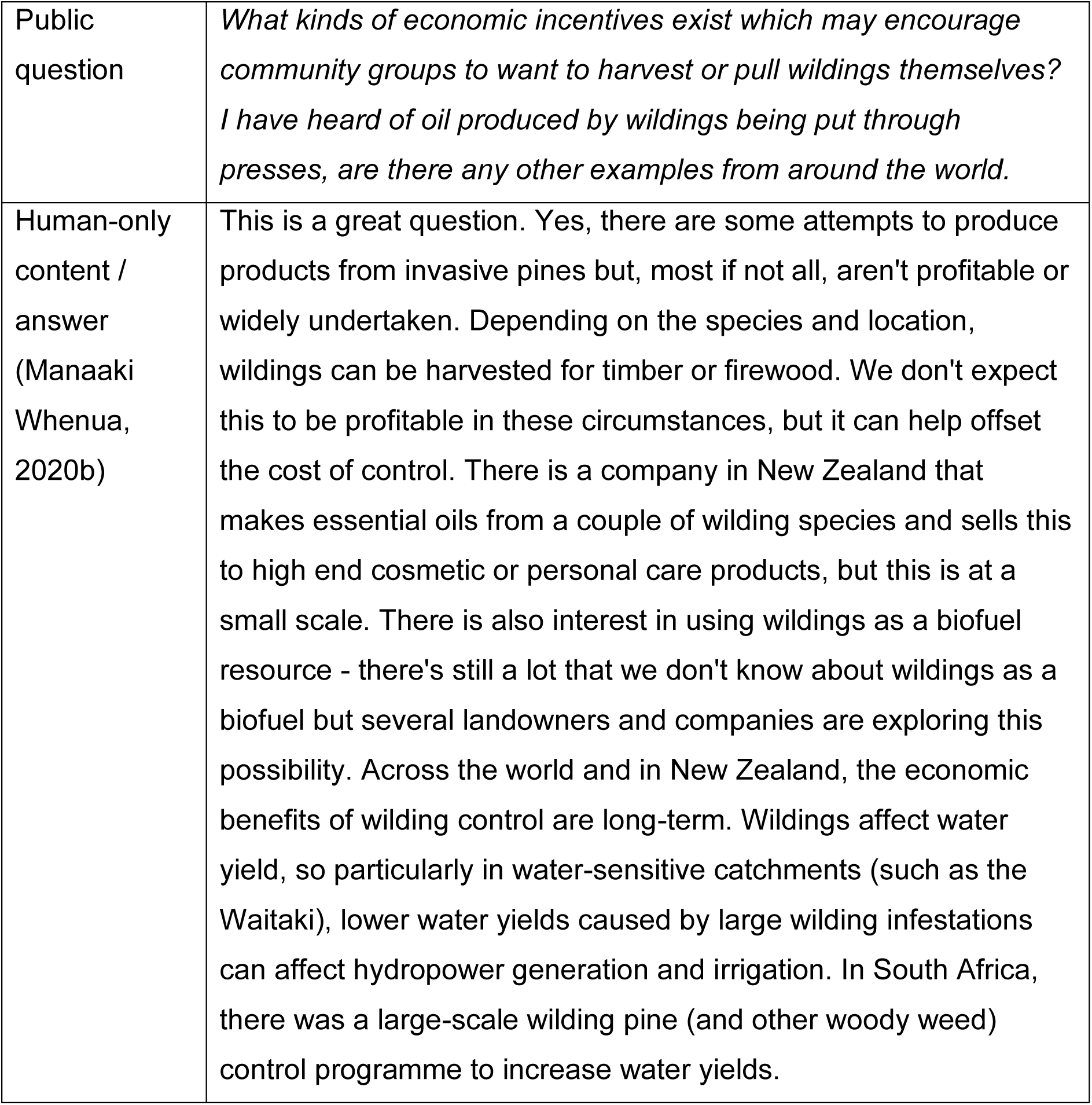
Human-only content Q&A Invasive Species.

| Public question | *What kinds of economic incentives exist which may encourage community groups to want to harvest or pull wildings themselves? I have heard of oil produced by wildings being put through presses, are there any other examples from around the world.* |
|---|---|
| Human-only content / answer (Manaaki Whenua, 2020b) | This is a great question. Yes, there are some attempts to produce products from invasive pines but, most if not all, aren't profitable or widely undertaken. Depending on the species and location, wildings can be harvested for timber or firewood. We don't expect this to be profitable in these circumstances, but it can help offset the cost of control. There is a company in New Zealand that makes essential oils from a couple of wilding species and sells this to high end cosmetic or personal care products, but this is at a small scale. There is also interest in using wildings as a biofuel resource - there's still a lot that we don't know about wildings as a biofuel but several landowners and companies are exploring this possibility. Across the world and in New Zealand, the economic benefits of wilding control are long-term. Wildings affect water yield, so particularly in water-sensitive catchments (such as the Waitaki), lower water yields caused by large wilding infestations can affect hydropower generation and irrigation. In South Africa, there was a large-scale wilding pine (and other woody weed) control programme to increase water yields. |

**Table S2.** GenAI Content Q&A Invasive Species.

|  |
|---|
| That's a great question. There are a few economic incentives that could encourage community groups to want to harvest or pull wildings themselves.<br>One incentive is the potential for financial compensation. In some cases, government agencies or private landowners will pay community groups to remove wilding conifers from their land. This can be a significant source of income for community groups, especially in rural areas where employment opportunities are limited.<br>Another incentive is the potential for environmental benefits. Removing wilding conifers can help to improve water quality, reduce erosion, and restore native habitats. This can have a positive impact on the local ecosystem, and it can also attract tourists and boost the local economy.<br>Finally, there is the potential for social benefits. Removing wilding conifers can be a great way for community groups to get involved in their local environment and to work together towards a common goal. This can build community spirit and pride, and it can also help to educate people about the importance of conservation.<br>As for your question about oil produced by wildings, there are a few examples of this from around the world. In New Zealand, a company called Wilding Conifer Oil is extracting oil from wilding pines and using it to make a variety of products, including biofuel, cosmetics, and pet food. In the United States, a company called Wildwood Oil is doing the same thing with wilding pines in Oregon.<br>These are just a few examples of the economic incentives that could encourage community groups to want to harvest or pull wildings themselves. By providing financial compensation, environmental benefits, and social benefits, we can help to make wilding conifer management a more attractive option for community groups.<br>In addition to the economic incentives I mentioned, there are also a few other things that could be done to encourage community groups to get involved in wilding conifer management. One is to provide training and education about wilding conifers and their impacts. This would help community groups to understand the importance of wilding conifer management and to develop effective management strategies.<br>Another thing that could be done is to provide access to equipment and resources. This would make it easier for community groups to remove wilding conifers from their land. Finally, it is important to build relationships with community groups and to listen to their concerns. This will help to ensure that wilding conifer management strategies are developed that are appropriate for local needs and conditions.<br>By taking these steps, we can help to make wilding conifer management a more effective and sustainable solution to this serious problem. |

**Figure S3.**
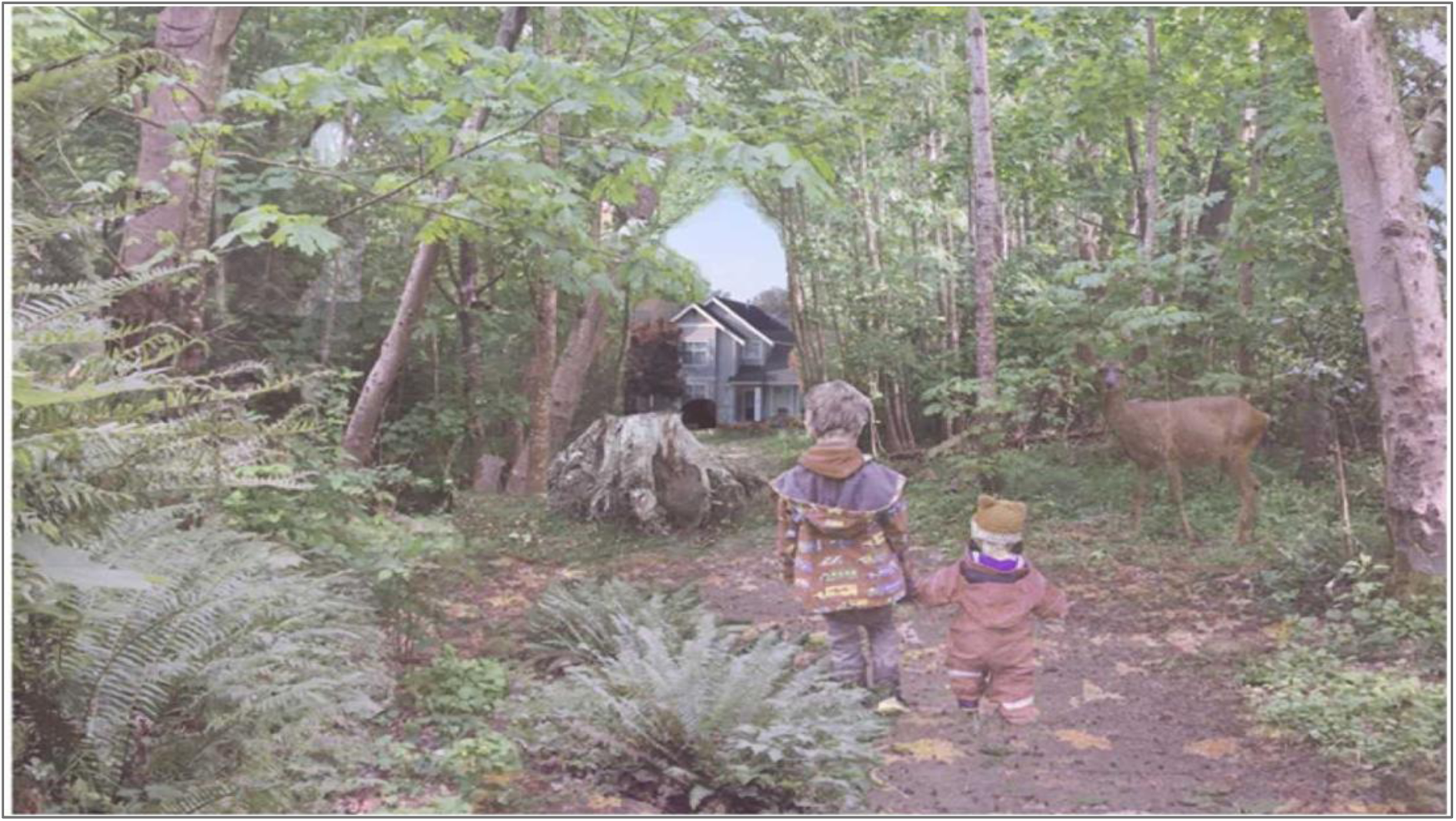
Barron et al. 2023 image of habitat restoration

**Figure S4.**
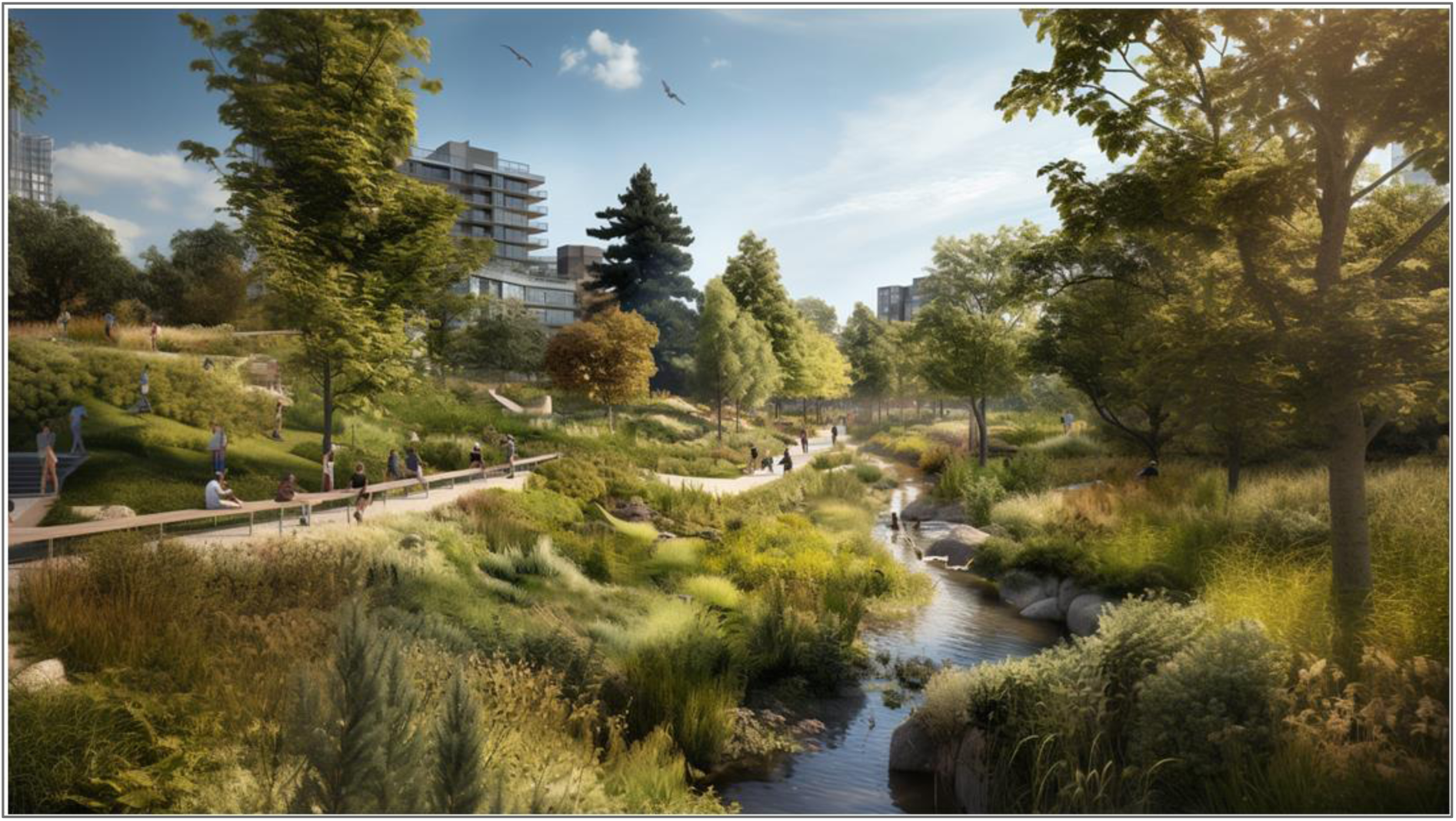
GenAI image of habitat restoration

### Supplementary Methods

#### Prompts used to generate Wetland Planting / Restoration Guide content

###### Prompts

- Hello, I would like to write a guidance document for the public in New Zealand about wetland creation and planting. The audience is varied, ranging from farmers, to conservation groups, to schools. I have some titles for subsections, please can you write the relevant text for each section? I will give you an approximate word limit for each section.
- PREPARE A PLANTING PLAN (128 words)
- Thank you. Now, please prepare a section on “weeds and pests” in only 35 words
- Thank you, now a section “timing” please in around 50 words. Please note that this section refers to both the season of planting and also any phasing of planting over several years.
- Thank you. Now please a section that highlights the steps for planting. This section is titled “Planting” with around 200 words
- Thank you. Now please could you help recommend specific species to plant in specific wetland zones? Please bear in mind that the context of this wetland planting is in New Zealand, and please follow your own advice provided in the sections above. First, what species should we plant in the shallow water planted zones (0.2 - 0.4 m depth).
- Thanks, how about the wet margins zone (less than 0.2 m depth)
- Thanks, how about on the wetland embankments?
- Thank you, and finally how about the surrounding riparian areas adjacent to the wetland itself

###### ChatGPT responses that are edited for formatting (changes noted by italics in square brackets)

[*Title: Prepare a Planting Plan*]

[*A planting plan is a crucial component of wetland creation, as it helps ensure the success and sustainability of the project. By following these steps, you can develop an effective planting plan:*]

1. [*Site Assessment:*] Begin by conducting a thorough site assessment. Identify the wetland’s type, soil conditions, hydrology, and existing vegetation. Consider any potential challenges or opportunities.
2. [*Select Native Species:*] Choose indigenous plant species that are well-adapted to the local climate and soil conditions. Native plants provide better habitat for wildlife, support biodiversity, and enhance ecosystem resilience.
3. [*Plant Diversity:*] Aim for a diverse mix of plant species to create a balanced ecosystem. Incorporate various types of grasses, sedges, rushes, wildflowers, and shrubs.
4. [*Seasonal Considerations:*] Plan for year-round interest and ecological benefits. Select plants that bloom at different times, providing food and shelter for wildlife throughout the year.
5. [*Planting Design:*] Lay out the planting areas and patterns, considering factors like water depth, sunlight exposure, and species’ growth requirements.
6. [*Maintenance Plan: Develop a maintenance schedule, including weed control, watering, and monitoring plant growth and establishment.*]

Remember to involve stakeholders [*,including farmers, conservation groups, and schools, in the planning process. Their active participation will foster a sense of ownership and community commitment to the wetland project.*]

Successful planting is essential for establishing a thriving wetland ecosystem. Follow these steps to ensure a smooth and effective planting process:

1. Site Preparation: Clear the planting area of weeds and non-native vegetation. Ensure the soil is moist but not waterlogged.
2. Handling Plants: Handle plants with care to protect their roots and delicate structures. Keep the roots moist and shaded during transportation.
3. Planting Technique: Dig appropriate-sized holes for each plant, ensuring they fit snugly without bending the roots. Avoid planting too deep or too shallow.
4. Spacing and Density: Follow the planting plan’s guidelines for spacing between plants and overall density. This promotes healthy growth and prevents overcrowding.
5. Watering: Water the plants immediately after planting, and continue regular watering during dry periods. Avoid overwatering, as excessive moisture can harm the plants.
6. Mulching: Apply a layer of organic mulch around the plants to retain moisture, suppress weeds, and protect against temperature fluctuations.
7. Maintenance: Regularly inspect the planted area for weeds, pests, and signs of stress in the plants. Address any issues promptly to ensure the successful establishment of the wetland.
8. [*Community Engagement: Involve local farmers, conservation groups, and schools in the planting process. Organize planting events to raise awareness and foster a sense of responsibility towards the wetland’s growth and protection.*]
9. [*Phased Planting: If planting over several years, prioritize the initial establishment of native species, then introduce additional plants in subsequent phases. This approach provides earlier habitat benefits while ensuring a diverse and resilient wetland ecosystem over time.*]

[*By following these planting steps and involving diverse stakeholders, you contribute to the creation of a vibrant wetland that supports biodiversity, water quality, and environmental sustainability in New Zealand.*]

##### 2. Prompts used to Respond to a Query from the Public

###### Initial prompt

Are you familiar with the youtube video “LINKOnline Webinar: New Zealand’s Worst Weed Problem” found here: https://www.youtube.com/watch?v=h1Ie9UrRlVY?

###### *Bard* response

Yes, I am familiar with the YouTube video “LINKOnline Webinar: New Zealand’s Worst Weed Problem”. It is a 50-minute webinar that was presented by the New Zealand Wilding Conifer Group in 2021. The webinar provides an overview of the wilding conifer problem in New Zealand, including the causes, consequences, and management interventions that are being used to address it. The webinar also highlights the importance of research and collaboration in developing effective wilding conifer management strategies…[response goes on to provide a further summary]

###### Follow-up prompt

Pretend you are one of the presenters. How would you answer this question from the audience: What kinds of economic incentives exist which may encourage community groups to want to harvest or pull wildings themselves? I have heard of oil produced by wildings being put through presses, are there any other examples from around the world.

##### Prompts used to generate park scenario images

As per the supplemental material from Barron et al. (2023) the text description of this scene is as follows:

“Habitat

*Neighbourhood residents express a desire for a sense of regional identity and chooses to replant with locally endemic species that will attract and sustain wildlife. Areas not yet developed are preserved and enhanced as wildlife areas, making a larger area of potential growing space (area with sufficient soil volume to support trees) than the previous two scenarios. Parks add habitat for local fauna with dense plantings with many structural shrub layers. Two major streets are closed to traffic and replaced with wildlife corridors, creating habitat connectivity for fauna.”*

*Midjourney* is a GenAI tool that allows user to input a text description to output images. We modified the text-description above to be able to prompt *Midjourney* as per the prompt below:

/imagine: no water, landscape, urban park, canada, Neighbourhood residents express a desire for a sense of regional identity and chooses to replant with locally endemic species that will attract and sustain wildlife. Areas not yet developed are preserved and enhanced as wildlife areas, making a larger area of potential growing space (area with sufficient soil volume to support trees). Parks add habitat for local fauna with dense plantings with many structural shrub layers. Two major streets are closed to traffic and replaced with wildlife corridors, creating habitat connectivity for fauna, ar 16:9.

Note that *Midjourney* prompts are initiated with the command [/imagine] followed by the description. We provided some additional context, that the image should not focus on bodies of water and that it should be reflective of a landscape of an urban park in Canada. We also specified the aspect ratio which allowed for us to compare more directly with the original human-only image without having to undertake cropping of the images. *Midjourney* produces four images at a time following a prompt and those four initial images are provided in Figure S5 below. Following from these four images we selected one image to be used in the survey to serve as a comparison to the human-only image from Barron et al. (2023).

The figure presented below (Figure S5) displays the four images resulting from prompting *Midjourney* using the following prompt modified from Barron et al. (2023). The modifications from the original Barron et al. (2023) text are presented in bold font:

**[/imagine: no water, landscape, urban park, canada,** Neighbourhood residents express a desire for a sense of regional identity and chooses to replant with locally endemic species that will attract and sustain wildlife. Areas not yet developed are preserved and enhanced as wildlife areas, making a larger area of potential growing space (area with sufficient soil volume to support trees). Parks add habitat for local fauna with dense plantings with many structural shrub layers. Two major streets are closed to traffic and replaced with wildlife corridors, creating habitat connectivity for fauna**, ar 16:9.]**

**Figure S5.**
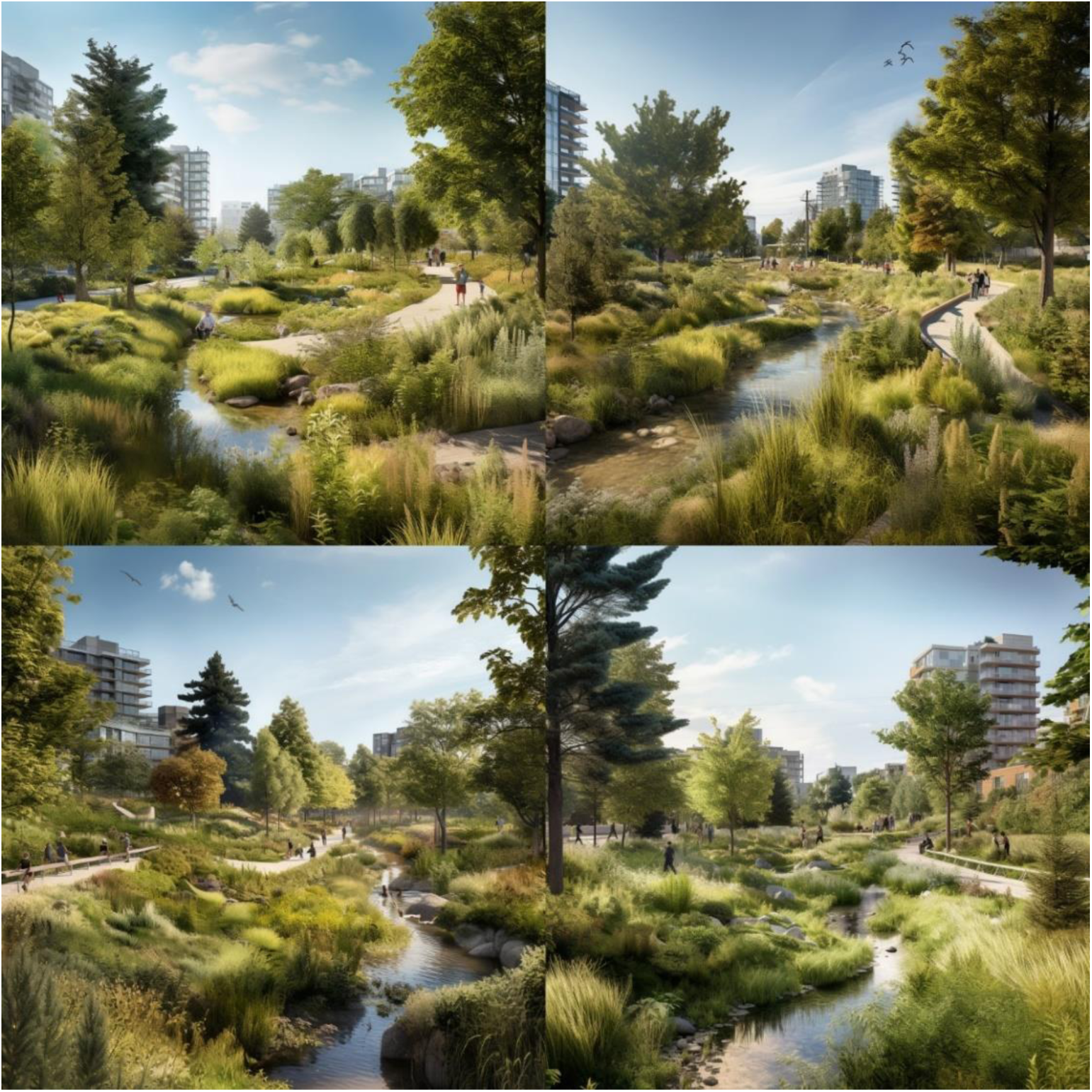
Initial Images from Midjourney

